# SAFB associates with nascent RNAs to promote gene expression in mouse embryonic stem cells

**DOI:** 10.1101/2022.12.20.521195

**Authors:** Rachel E. Cherney, Quinn E. Eberhard, Christine A. Mills, Alessandro Porrello, Zhiyue Zhang, David White, Laura E. Herring, J. Mauro Calabrese

## Abstract

Scaffold Attachment Factor B (SAFB) is a conserved RNA Binding Protein (RBP) that is essential for early mammalian development. However, the RNAs that associate with SAFB in mouse embryonic stem cells have not been characterized. Here, we addressed this unknown using RNA-seq and SAFB RNA immunoprecipitation followed by RNA-seq (RIP-seq) in wild-type ESCs and in ESCs in which SAFB and SAFB2 were knocked out. SAFB predominantly associated with introns of protein-coding genes through purine-rich motifs. The transcript most enriched in SAFB association was the lncRNA *Malat1*, which also contains a purine-rich region in its 5*′* end. Knockout of SAFB/2 led to down- and upregulation of approximately 1,000 genes associated with multiple biological processes, including genes that are regulated by Polycomb and genes involved in apoptosis, cell division, and cell migration. The spliced and nascent transcripts of many downregulated genes associated with high levels of SAFB in wild-type cells, implying that SAFB binding promotes their expression. Reintroduction of SAFB into double-knockout cells restored gene expression towards wild-type levels, an effect that was again observable at the level of spliced and nascent transcripts. Proteomics analysis revealed a significant enrichment of nuclear speckle-associated and RS-domain containing proteins among SAFB interactors. Our findings suggest that among other potential functions in mouse embryonic stem cells, SAFB promotes the expression of a subset of genes through its ability to bind purine regions in nascent RNA.

## Intro

SAFB is a chromatin-associated RNA-binding protein that has been implicated in multiple molecular processes including the regulation of transcription, splicing, and chromatin structure (Garee and Oesterreich 2010; Norman et al. 2016). SAFB was originally identified as a chromatin-associated protein that could bind specific DNA sequences *in vitro* and repress transcription from an estrogen-responsive promoter upon its transient over-expression (Renz and Fackelmayer 1996; Oesterreich et al. 1997). Whether SAFB associates with DNA *in vivo* remains an open question. However, SAFB contains an RNA Recognition Motif (RRM) and an RGG/RG motif, both of which are known RNA-binding domains (Corley et al. 2020). Moreover, *in vivo*, SAFB has been shown to co-localize with other RNA-binding proteins, co-purify with mRNA, and bind RNA sequences that are purine rich, solidifying its role as bona fide RNA-binding protein (Nayler et al. 1998; Weighardt et al. 1999; Arao et al. 2000; Baltz et al. 2012; Castello et al. 2012; Hong et al. 2015; Rivers et al. 2015). SAFB has also been implicated in regulating the response to heat shock in human cells, where along with other RNA-binding proteins, it becomes enriched in nuclear condensates centered around specific satellite RNAs (Aly et al. 2019). Additionally, SAFB plays a role in the nuclear retention of unspliced RNAs (Ron and Ulitsky 2022), and has been shown to be important for the maintenance of heterochromatin in mouse and human cells, possibly through its ability to associate with specific RNAs (Huo et al. 2020; McCarthy et al. 2021).

Consistent with the involvement of SAFB in more than one fundamental cellular process, SAFB is required for proper mouse development. While SAFB null embryos can survive to term, most die shortly after birth, and in surviving animals, pleiotropic abnormalities that include defects in the endocrine system persist throughout life (Ivanova et al. 2005). SAFB also has a paralogue, SAFB2, which shares an overall domain structure with and is similar in sequence to SAFB but whose loss does not cause embryonic lethality (Jiang et al. 2015). Still, SAFB2 is highly expressed in the male reproductive tract, where it may play important roles in endocrine signaling (Jiang et al. 2015). SAFB2 is also important the processing of several miRNAs (Hutter et al. 2020).

Despite the critical roles of SAFB in development, its RNA targets have not been profiled in early embryonic tissues or cell lines derived from them. Herein, we describe the generation of SAFB and SAFB2 double-knockout (DKO) mouse embryonic stem cells (ESCs) and data from subsequent RNA immunoprecipitation (RIP) and RNA-Seq experiments that identify the RNA targets that associate with SAFB in ESCs. We found that SAFB associates predominantly with intronic regions of protein coding genes. Genes whose expression levels were reduced after SAFB/2 knockout associated more robustly with SAFB in wild-type cells than genes whose expression levels increased, suggesting that at the steady-state, SAFB can promote the expression of a subset of genes through its ability to associate with nascent RNA. By proteomics, we found that SAFB associates with many proteins found in speckles – proteinaceous nuclear condensates that have been shown to promote transcription of nearby genes (Kim et al. 2020; Zhang et al. 2020). Our data provide insights into the molecular functions of SAFB in the early stages of embryogenesis and suggest a model whereby the aggregate protein-binding profile of a nascent RNA may impact gene expression by modulating the proximity of its corresponding host gene to different classes of nuclear condensate (Sabari et al. 2020).

## Results

### Generation and validation of SAFB/2 double knockout mouse embryonic stem cells

We generated SAFB and SAFB2 DKO ESCs using CRISPR-mediated genome editing in an F1-hybrid male ESC line that we previously engineered to express the *Xist* long noncoding RNA (lncRNA) under the control of a doxycycline-inducible promoter from the *Rosa26* safe-harbor locus located on the B6-derived copy of chromosome 6 (Trotman et al. 2020). Single guide RNA sequences flanking the SAFB/2 locus on chromosome 17 were designed using CRISPOR (Concordet and Haeussler 2018), cloned into pX330 (Cong et al. 2013), and transfected into ESCs. After a short pulse of puromycin, ESCs were plated at low density onto feeder cells to enable picking of individual colonies. Two individual colonies that genotyped as double-knockouts by DNA PCR, reverse-transcription coupled PCR (RT-PCR), and western blot were selected for our study below (DKO4 and DKO13; Figure 1).

**Figure 1.**
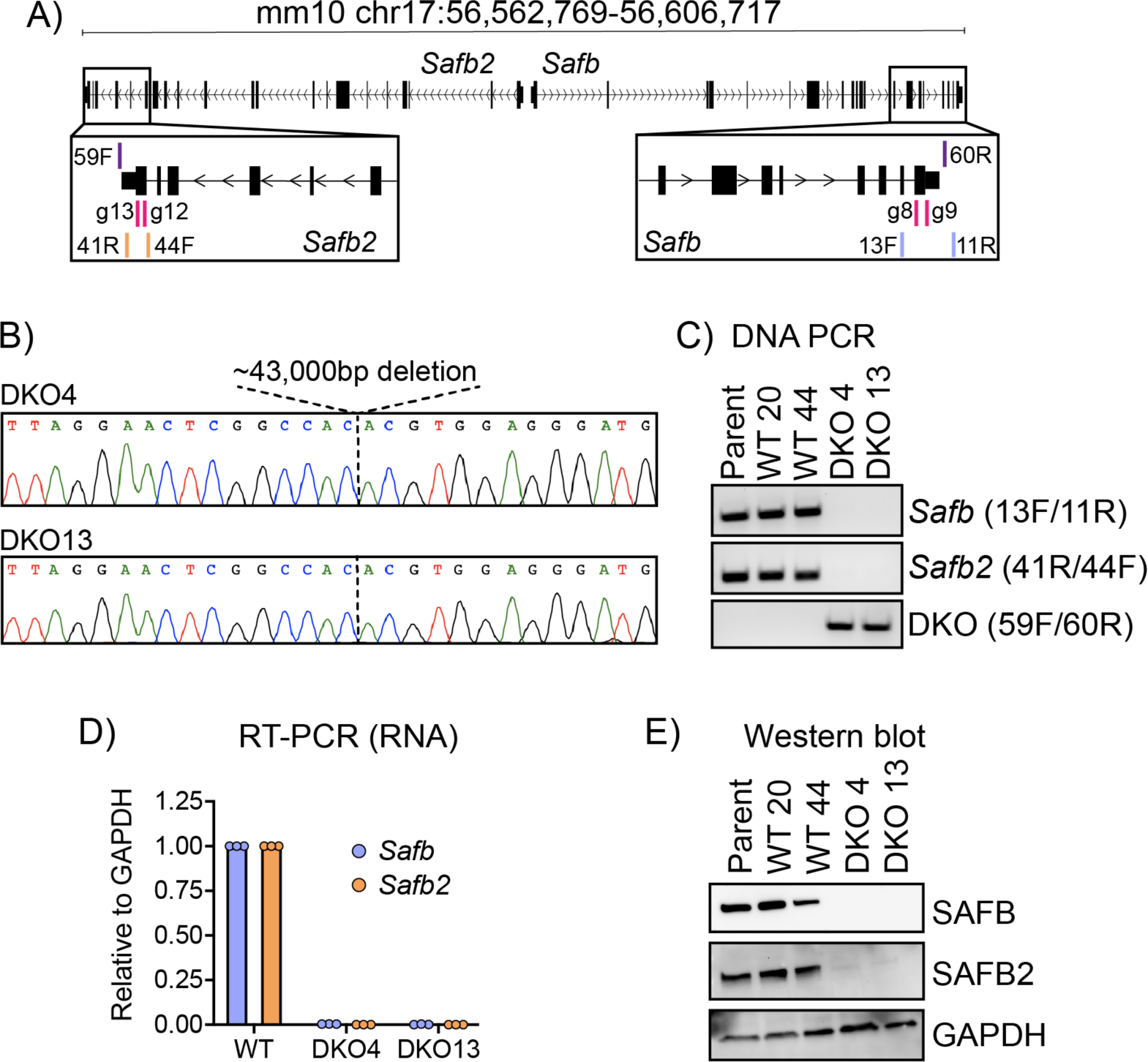
Generation and validation of SAFB/2 double knockout mouse embryonic stem cells. **(A)** Schematic of *Safb* and *Safb2* mm10 genomic locus with location of sgRNAs (g8-g12) and genotyping primers (13F/11R; 41R/44F; 59F/60R). **(B)** Sanger sequencing traces through the deleted region in the two clonal knockout lines used in this study, DKO4 and DKO13. **(C)** PCR genotyping in wild-type (WT) and double knock out (DKO) ESCs. **(D)** qPCR for RNA detection of *Safb* and *Safb2* mRNA levels (shown relative to GAPDH in wild-type cells). Dots represent technical triplicate measurements. **(E)** Western blots showing levels of SAFB and SAFB2 in wild-type and DKO cells.

### SAFB associates predominantly with intronic regions of protein-coding genes in mouse embryonic stem cells

To identify RNAs that associate with SAFB in ESCs, we used formaldehyde-based RNA immunoprecipitation (RIP; (Raab et al. 2019)) and a polyclonal rabbit antibody raised against the C-terminal region of SAFB, which we purchased from Bethyl Laboratories (product # A300-812A). RIPs were performed in five separate ESC lines: the wild-type, *Xist*-expressing ESC line from (Trotman et al. 2020) (“Parent”), two clonal ESC lines that underwent the pX330 transfection and colony picking but remained wild-type for SAFB (WT20 and WT44), and the two DKO ESC lines described in Figure 1. Proportionally equal volumes of RNA prepared from each RIP were then subjected to cDNA library preparation and sequencing as in (Schertzer et al. 2019a). We performed all RNA collection experiments under the doxycycline-induced condition, in which *Xist* is expressed from a single copy of chromosome 6 (Trotman et al. 2020).

To identify regions of RNA enriched in their association with SAFB (i.e. “SAFB peaks”), we used a strand-specific implementation of MACS2 and a combination of DESeq2 and empirical filtering of regions based on their signal in wild-type ESCs relative to SAFB/2 DKO controls (Zhang et al. 2008; Feng et al. 2012; Love et al. 2014). Specifically, we required that enriched regions (1) were represented by at least five reads in at least two of the three WT ESC lines profiled, (2) were ascribed a p-value of <0.05 by DESeq2 when comparing signal between SAFB/2 DKO ESCs and wild-type ESCs, and (3) had an average reads-per-million (RPM) signal of at least two-fold less in SAFB/2 DKO ESCs compared to wild-type ESCs. This yielded 32,354 regions that were potentially enriched in their association with SAFB in wild-type ESCs. As an additional filter, in DKO13 ESCs, we stably expressed SAFB and nuclear-localized GFP cDNAs tagged at their C-termini with tandem FLAG and V5 epitopes (Figure 2A) and performed RIP-Seq using an anti-FLAG antibody. Out of the 32,354 potential peak regions identified above, 23,853 regions had a total of at least 5 reads in the SAFB-FLAG and GFP-FLAG datasets, and 94% of these regions (22,497) had a higher signal in the SAFB-FLAG dataset compared to the GFP-FLAG negative control, supporting the high fidelity of our endogenous SAFB RIP data as well as our peak calling approach. The 1,356 regions with higher signal in the GFP-FLAG compared to the SAFB-FLAG RIP were dropped from further analysis, yielding a total of 30,998 regions that we henceforth define as SAFB-associated peaks (Table S1).

**Figure 2.**
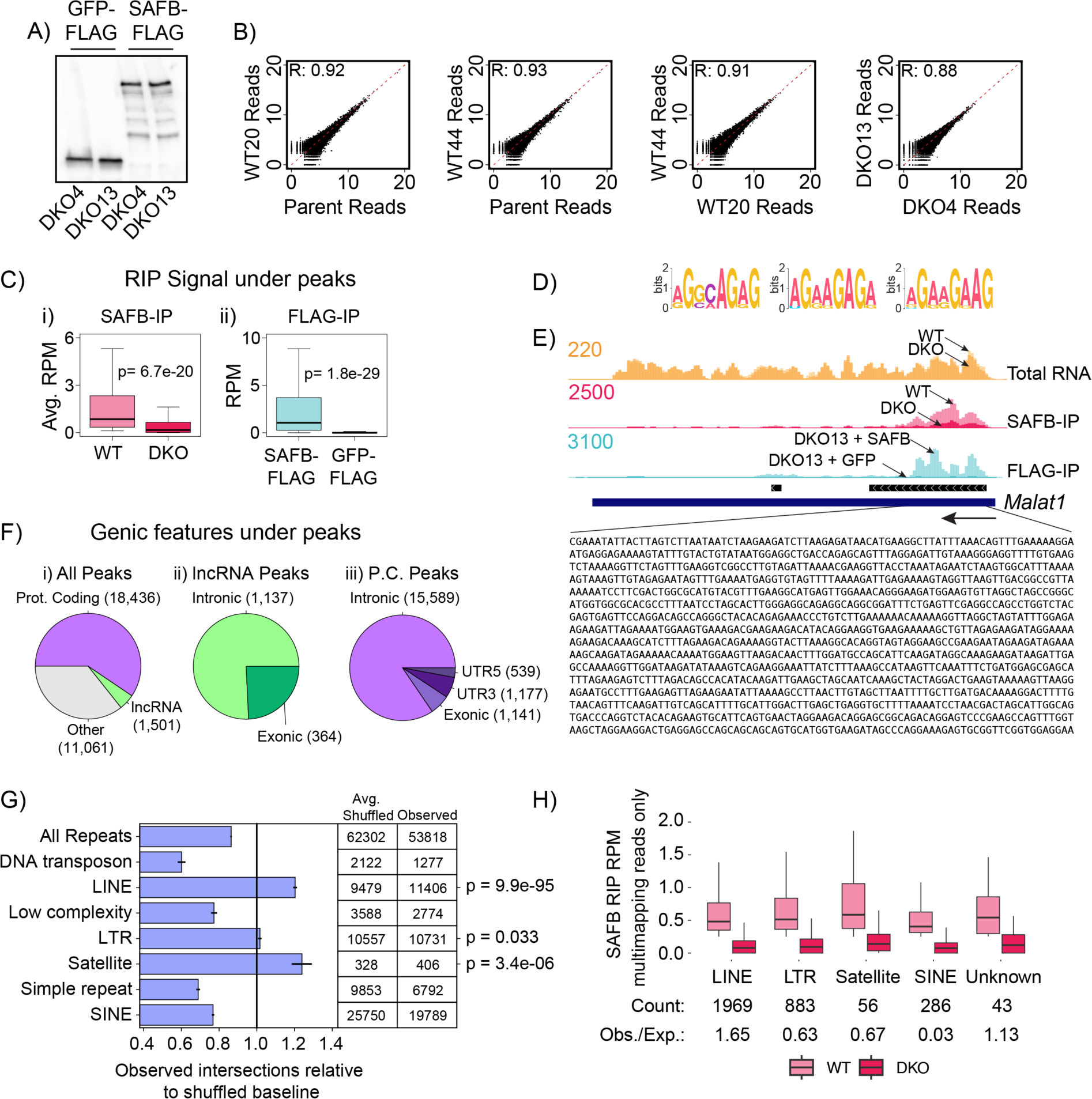
SAFB associates predominantly with intronic regions of protein-coding genes in mouse embryonic stem cells. **(A)** αFLAG western blot of GFP and SAFB rescue in DKO ESCs. **(B)** Scatter plots showing correlation of RIP-Seq replicate data under each SAFB peak for each genotype. **(C)** Reads-per-million (RPM)-normalized RIP signal under SAFB peaks in (i) wild-type and DKO ESCs (αSAFB RIP) and (ii) in SAFB and GFP rescue ESCs (αFLAG RIP). **(D)** Top three motifs derived from analysis of sequence under SAFB peaks. **(E)** RNA Input and RIP-Seq wiggle density tracks at the *Malat1* locus. Black rectangles under wiggle tracks denote location of SAFB peaks. **(F)** Genic features that overlap SAFB peaks. **(G)** Intersection of mm10 annotated repeat elements and SAFB peaks. p-values = (1 - [cumulative distribution function]) of the distribution of intersections observed from 1,000 shuffles sets of SAFB peaks. **(H)** Intersection of repeat-masked elements and multimapping reads from SAFB RIPs in WT and DKO ESCs. “Count”, number of elements in each class that passed threshold for inclusion. “Obs./Exp”, ratio of observed vs. expected genomic space occupied by each class of repeat in the list of elements passing filter (Table S2). Classes of repeat that were represented by less than 25 elements in our filtered list were not plotted.

The signal patterns under each peak were highly reproducible between replicate RIP experiments. Comparing replicates between wild-type samples yielded an average Pearson’s r value of 0.92 and comparing replicates between DKO samples yielded a Pearson’s r value of 0.88 (Figure 2B). Boxplots of RIP signal under each peak in the different genotypes demonstrate the SAFB dependence of signal under SAFB peaks (Figure 2C). Wiggle density tracks of individual and pooled replicates can be viewed on the UCSC genome browser ((Lee et al. 2022); https://genome.ucsc.edu/s/recherney/Cherney_Safb_2022). Consistent with prior iCLIP studies of SAFB in human SH-SY5Y and MCF-7 cells, we identified GA-rich sequence motifs associated with the most-strongly enriched SAFB peaks (Figure 2D; (Hong et al. 2015; Rivers et al. 2015)). An image of SAFB RIP data over *Malat1*, the gene that exhibited the highest association with SAFB in our datasets, is shown in Figure 2E. *Malat1* has previously been shown to interact with SAFB in human cells (Hong et al. 2015; Spiniello et al. 2018). Moreover, the region within in *Malat1* that is most strongly associated with SAFB is visibly enriched in GA nucleotides, consistent with the association being driven by direct interactions (Figure 2E).

We next determined the location of SAFB peaks relative to the GENCODE vM25 “basic” gene annotation set (Frankish et al. 2019). We found that 18,436 SAFB peaks overlapped with transcripts originating from protein-coding genes and 1,501 peaks with transcripts originating from long noncoding RNAs (lncRNAs; Table S1). An additional 6,004 peaks were located within 10 kilobases (kb) of an annotated transcript in the GENCODE vM25 “comprehensive” gene annotation set. The majority of gene-overlapping peaks fell within intronic regions, for the set of peaks that overlapped both protein-coding and lncRNA transcripts (Figure 2F).

SAFB has previously been shown to associate with specific classes of repetitive elements, most notably those derived from satellite repeats and LINEs (Aly et al. 2019; Huo et al. 2020). Therefore, we determined the extent to which SAFB peaks were enriched in overlapping repetitive elements relative to sets of locally shuffled control peaks. Consistent with prior works, the strongest enrichments were observed over LINE- and satellite-derived elements (Figure 2G; Table S1). Next, because our set of SAFB peaks were defined using uniquely mapped, non-repetitive reads, we performed a repeat analysis exclusively with multimapping reads, using the TElocal package to assign fractional read counts to repeat-derived elements in the mm10 build of the genome (Jin et al. 2015). This analysis uncovered a set of 6162 repeat-derived elements that were detected above a minimal expression threshold and whose RPM-normalized expression values were, on average, at least 2-fold higher in the WT compared to DKO RIP-Seq data (Figure 2H; Table S2). Twenty-six percent of the elements in this set (1588/6162) overlapped a gene annotation in GENCODE (Frankish et al. 2019). While essentially all classes of repeats were represented, we observed a significant enrichment for LINE-derived elements, and significant depletions for all other classes of repeat (Figure 2H and not shown; p <2e-16 for all differences, Chi-squared). We conclude that within the context of uniquely alignable regions as well as repetitive genic and intergenic loci, the SAFB protein associates with LINE-derived elements at higher-than-expected frequencies. Additionally, on a local scale, uniquely alignable peaks of SAFB also overlap satellite-derived elements with a higher-than-expected frequency.

### SAFB/2 loss alters expression of a subset of genes in mouse embryonic stem cells

We next assessed whether SAFB DKO ESCs exhibited significant gene expression changes relative to WT ESCs. For this analysis, we performed total RNA-seq on “input” RNA collected from the same crosslinked ESCs that we used to perform SAFB RIP above. Using DESeq2, we identified 992 genes whose expression was significantly different (padj<0.05) between our three WT and two DKO samples; 545 of these genes were downregulated in DKO compared to WT ESCs, and 447 were upregulated (Figure 3A, B; Table S3).

**Figure 3.**
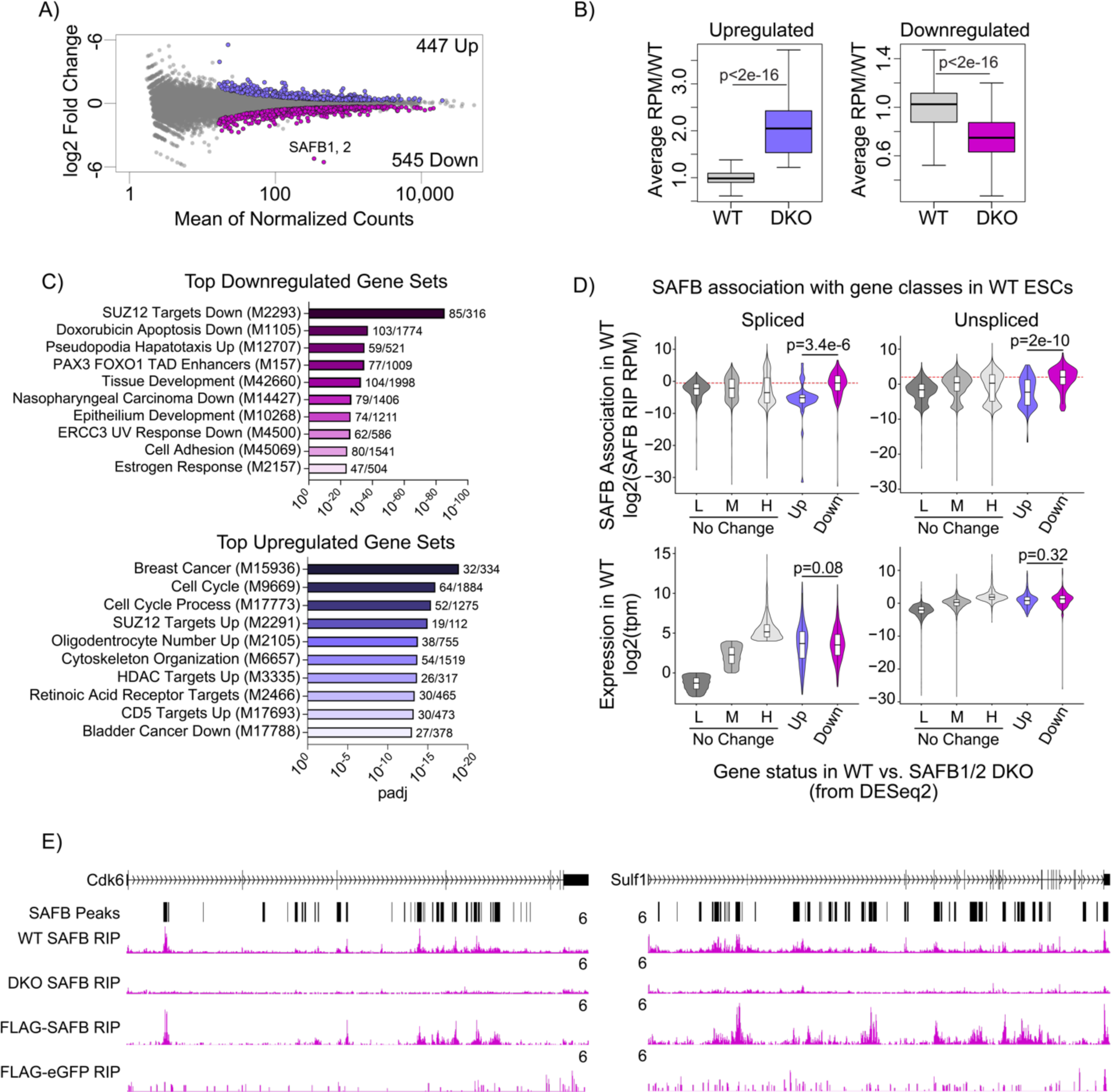
SAFB/2 loss alters expression of a subset of genes in mouse embryonic stem cells. **(A)** MA plot of differential gene expression between wild-type and DKO ESCs; purple dots, upregulated genes, magenta dots, downregulated genes. **(B)** Average RPM of up and down regulated genes in wild-type relative to DKO ESCs. p-values, paired t-test. **(C)** Top ten enriched gene sets (from MSigDB; (Subramanian et al. 2005; Liberzon et al. 2015)) associated with up- and downregulated genes. **(D)** SAFB association and overall expression levels of spliced and unspliced transcripts in wild-type ESCs, broken down by whether genes are upregulated (“Up”), downregulated (“Down”), or do not change in expression upon SAFB/2 DKO (“No Change”). The set of non-changing genes is further partitioned into three categories: those with low (0.125 -1 TPM; “L”), medium (>1 but <16 TPM; “M”),) and high (>16 TPM; “H”),) levels of expression. **(E)** Representative genes (*Cdk6* and *Sulf1*) that harbor intronic peaks of SAFB association and whose expression drops upon DKO and is restored upon reintroduction of SAFB cDNA.

Using Gene Set Enrichment Analysis and the Molecular Signatures Database (MSigDB; (Subramanian et al. 2005; Liberzon et al. 2015)), we examined the genes and pathways that were significantly associated with the differentially expressed genes. Specifically, we queried MSigDB’s “Hallmark Gene”, “Chemical and Genetic Perturbation”, and “GO Biological Process” gene set collections (Table S4). Among the genes that were downregulated in SAFB DKO cells, we identified several significant connections with gene sets involved in tissue development, cell adhesion, and the response to estrogen (Dutertre et al. 2010), among others (Figure 3C; Table S4). Intriguingly, the most significantly enriched gene set among the downregulated genes was “PASINI_SUZ12_TARGETS_DN”, a set of genes whose expression is significantly downregulated upon knockout of the Polycomb protein SUZ12 in ESCs (Figure 3C; (Pasini et al. 2007)).

Among the genes that were upregulated in DKO cells, enriched gene sets were also identified, although the enrichments were generally not as strong as those detected with the downregulated genes (Figure 3C; Table S4). However, we again identified a strong connection between SAFB/2 DKO and Polycomb-regulated genes; the fourth most significantly enriched gene set was “PASINI_SUZ12_TARGETS_UP”, the set of genes whose expression was significantly upregulated upon knockout of the Polycomb protein SUZ12 in mouse embryonic stem cells (Figure 3C; (Pasini et al. 2007)). Additional notable enrichments included genes involved in cycle progression, as well as targets of histone deacetylases (Heller et al. 2008) and the retinoic acid receptor (Delacroix et al. 2010), among others (Figure 3C; Table S4).

We next sought to determine whether the genes whose expression changed upon SAFB/2 loss exhibited evidence of SAFB association by RIP. Considering our prior observation that SAFB associates predominantly with introns, we took an approach that would let us quantify the extent of SAFB association with both spliced and unspliced RNAs. Briefly, starting with our wild-type and DKO RIP-Seq datasets, we extracted the reads that were aligned by STAR under each SAFB peak, and then used probabilistic alignment with kallisto to re-align those same reads to a version of the mouse transcriptome that contained one representative unspliced transcript for each GENCODE gene (Dobin et al. 2013; Bray et al. 2016; Frankish et al. 2021). We then used the difference of transcript per million (TPM) values in the wild-type and DKO RIP-Seq datasets to estimate the extent of SAFB association with each expressed transcript isoform. Strikingly, both the spliced and unspliced isoforms of the genes that were downregulated upon SAFB/2 DKO associated with significantly more SAFB than genes that were upregulated upon DKO or whose expression did not change (Figure 3D, upper panels; p<0.001 for both, paired t-test; Table S5). This difference could not be accounted for by differences in overall expression levels (Figure 3D; lower panels). Indeed, both the spliced and unspliced transcripts of the set of downregulated genes associated with significantly higher levels of SAFB than any other set of genes we examined, except the most highly expressed subset of spliced transcripts (Figure 3D; p<0.001 for all significant values, paired t-test; Table S5). These data imply that among the set of downregulated genes are many direct targets of SAFB, and that association with SAFB in these instances serves to promote overall gene expression.

Reintroduction of SAFB into SAFB/2 knockout cells restores gene expression defects in a manner dependent on the SAFB C-terminal domain

SAFB is comprised of multiple domains, most of which are important for its proper localization in mouse cells (Huo et al. 2020). We were intrigued by the final ∼300 amino acids of SAFB, an R/G-rich domain which is predicted to be intrinsically disordered and important for SAFB’s ability to interact with both proteins and RNA (Townson et al. 2004; Finn et al. 2016; Dosztanyi 2018; Meszaros et al. 2018; Corley et al. 2020; Huo et al. 2020). This same C-terminal domain is sufficient to mediate repression of a heterologous reporter gene when tethered to its promoter (Townson et al. 2004).

To study the role of the C-terminal domain in mediating the effects of SAFB on gene expression in ESCs, we introduced three separate expression constructs into DKO ESCs via piggyBac-mediated transgenesis. The first two constructs, described in Figure 2A above, constitutively express full-length SAFB and GFP cDNAs each tagged at their 3*′* ends with 3xFLAG and V5 epitopes and a nuclear-localization signal. A third construct in the same vector backbone expresses a mutant version of SAFB in which the C-terminal disordered domain has been deleted (Figure 4A, *Δ*DD3). Western blot confirmed expression of all three constructs and additionally demonstrated that *Δ*DD3 is more highly expressed than full-length SAFB (Figure 4B). RIP-qPCR using either *α*FLAG or *α*V5 antibodies demonstrated that the DD3 domain is required for SAFB association with target RNAs, consistent with expectations (Figure 4C; (Huo et al. 2020)).

**Figure 4.**
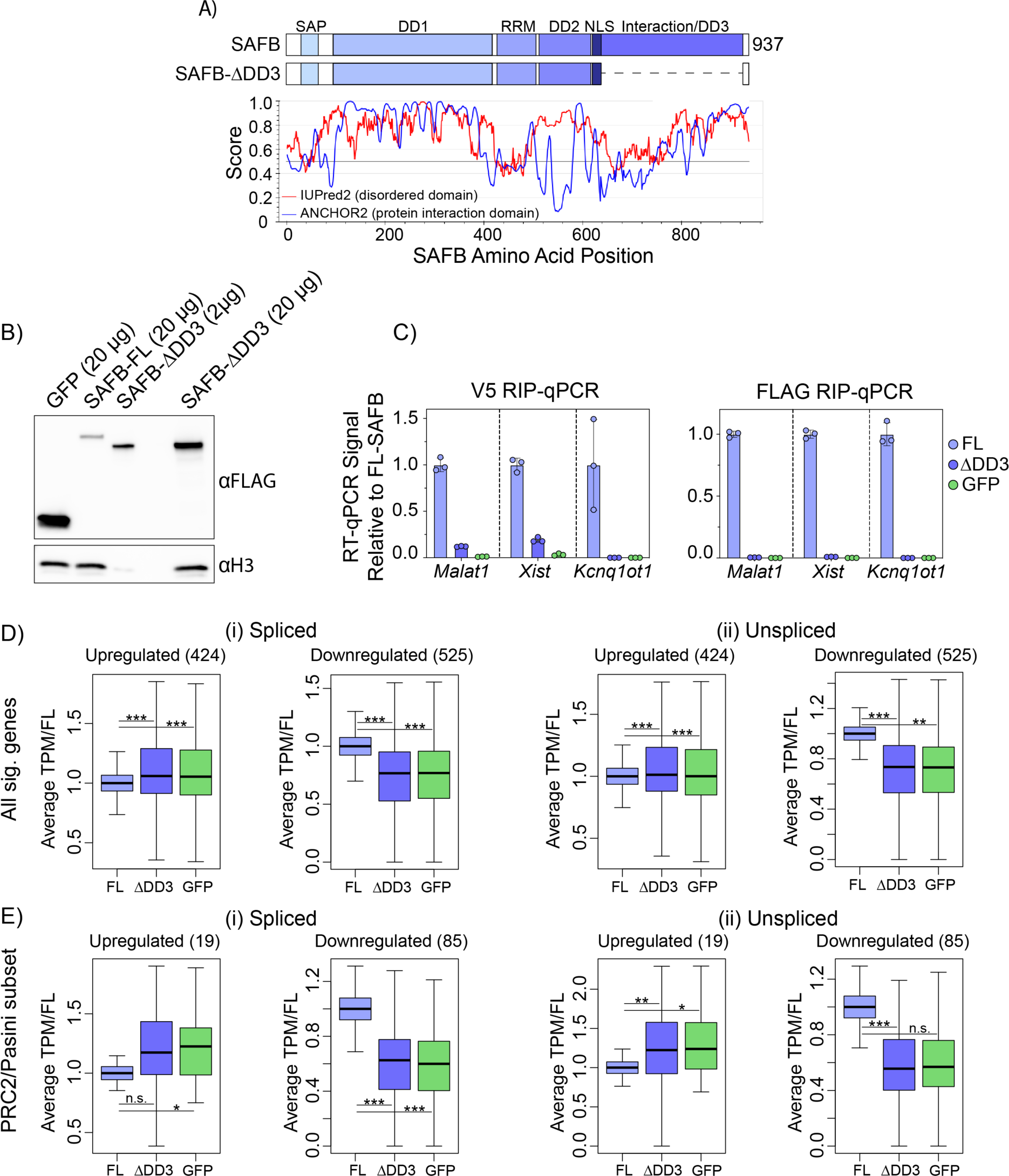
Reintroduction of SAFB into SAFB/2 knockout cells restores gene expression defects in a manner dependent on the SAFB C-terminal domain. **(A)** Protein domain diagram of SAFB (top) and ΔDD3 (bottom). IUPred2 disorder predictions below. **(B)** αFLAG western blot of GFP and SAFB rescue in DKO ESCs. **(C)** αV5 and αFLAG RIP-qPCR signal relative to FL-SAFB in FL-SAFB, ΔDD3, and GFP rescue cells. **(D,E)** Boxplots of average TPM in FL-SAFB, ΔDD3, and GFP rescue cells. *, **, ***, p-values <0.05, <0.01, and <0.001, respectively.

To determine whether reintroduction of SAFB restored gene expression defects in DKO ESCs, we performed RNA-Seq from biological duplicate preparations of RNA extracted from full-length SAFB, *Δ*DD3, and GFP-expressing DKO cells. Using kallisto to estimate the abundance of spliced and unspliced isoforms (Bray et al. 2016), we found that spliced and unspliced isoforms of transcripts produced from the genes that we had previously found to be significantly upregulated or downregulated upon SAFB/2 DKO returned closer to wild-type levels upon expression of full-length SAFB but not *Δ*DD3 or GFP (Figure 4D). The trends were numerically stronger for the set of downregulated genes compared those that were upregulated (Figure 4D). That the downregulated genes shifted more strongly towards wild-type levels upon reintroduction of SAFB is consistent with our prior observation that the nascent RNAs produced from many downregulated genes associate with high levels of SAFB (Figure 3D). Examining the subsets of genes that were up- and downregulated in DKO cells and similarly dysregulated upon SUZ12 knockout in ESCs, we observed analogous but numerically stronger trends (Figure 4E; Table S5). We conclude that many of the transcriptional changes that occur upon SAFB/2 DKO can be restored by reintroduction of SAFB into DKO cells, and that the restoration of these changes depends on the C-terminal domain of SAFB, if not other regions of the protein as well (Huo et al. 2020).

### The C-terminal region of SAFB interacts with RS-domain containing and speckle-associated proteins

SAFB has been shown to interact with several SR proteins through its C-terminal region, and is also found in nuclear speckles, which are nuclear condensates that harbor high levels of SR proteins and are associated with increased expression of surrounding genes (Nayler et al. 1998; Saitoh et al. 2004; Townson et al. 2004; Kim et al. 2020; Zhang et al. 2020). To determine whether SAFB interacts with SR proteins and speckle components in ESCs, we performed mass spectrometry-based proteomics on biological duplicate preparations of proteins immunoprecipitated by the FLAG antibody from formaldehyde-crosslinked extracts made from DKO cells expressing the same the full-length SAFB, *Δ*DD3, and GFP cDNAs described in Figure 4.

SAFB was originally identified along with another abundant RNA-binding protein called HNRNPU (or SAF-A), owing to their mutual presence in high salt extractions from HeLa nuclei and their ability to bind hydroxylapatite columns and S/MAR DNA elements *in vitro* (Romig et al. 1992; Renz and Fackelmayer 1996). Similar to that observed for SAFB above, HNRNPU has previously been implicated in promoting gene expression through its ability to associate with nascent, chromatin-associated RNA (Nozawa et al. 2017). We reasoned that comparing the proteins that are associated with SAFB and HNRNPU in ESCs might shed light on whether they promote gene expression using shared or different mechanisms. Therefore, in our wild-type parent ESC line, we expressed a FLAG-tagged cDNA of HNRNPU as well as a version of HNRNPU lacking a 154 amino acid-long, R/G-rich region at its C-terminus (Figure 5A, B; *Δ*RGG). We then performed mass spectrometry-based proteomics in technical duplicate from a single biological replicate of proteins immunoprecipitated by the FLAG antibody from formaldehyde-crosslinked extracts of HNRNPU and *Δ*RGG-expressing wild-type ESCs.

**Figure 5.**
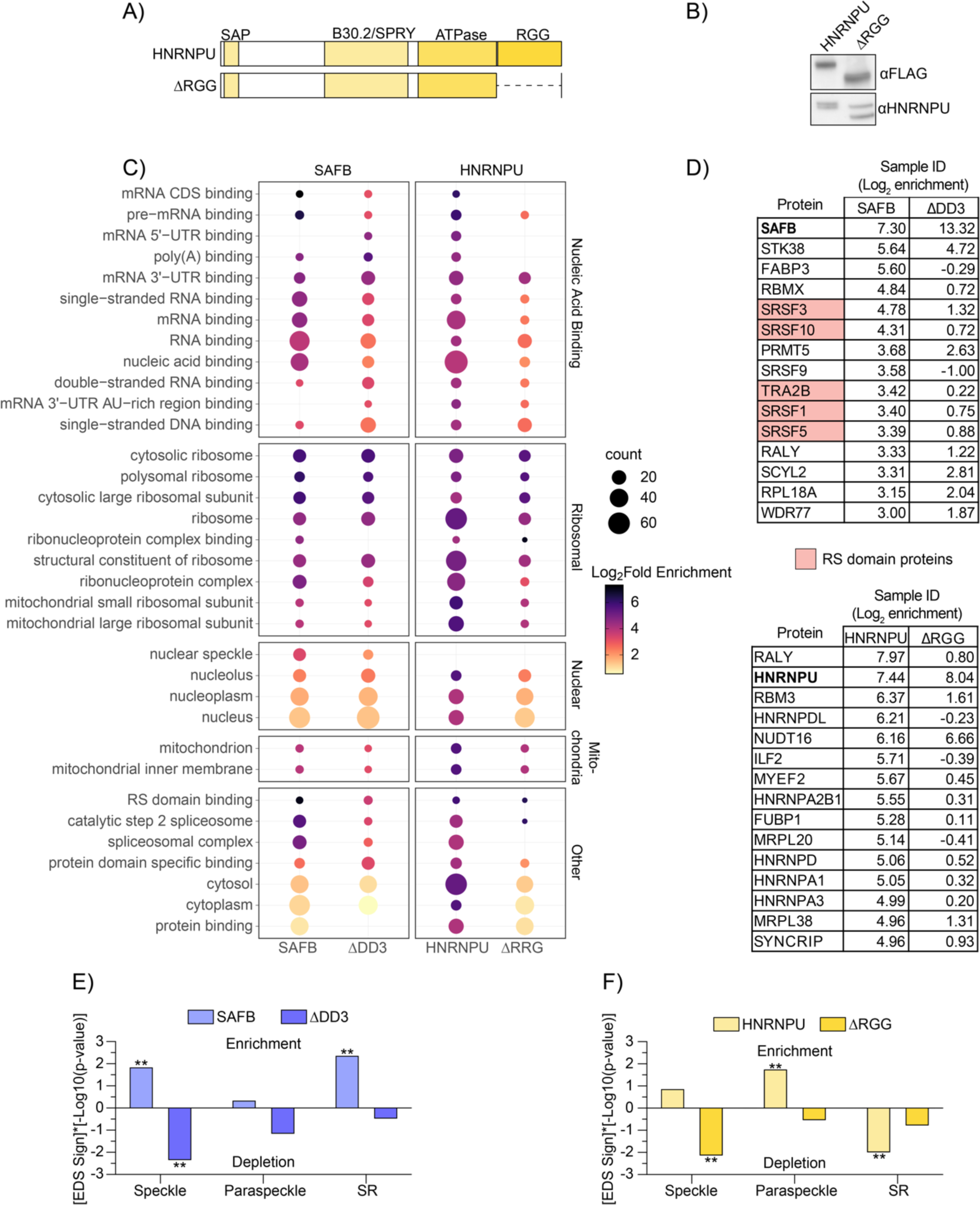
The C-terminal region of SAFB interacts with RS-domain containing and speckle-associated proteins. **(A)** Protein domain diagram of HNRNPU (top) and ΔRGG (bottom). **(B)** αFLAG and αHNRNPU western blot analyses of HNRNPU and ΔRGG-expressing cells. In the blot probed with αHNRNPU (lower panel), both endogenous HNRNPU and FLAG-tagged HNRNPU are visible. **(C)** Top GO terms from SAFB and HNRNPU IPs. **(D)** Top 15 most-enriched proteins in (i) SAFB and (ii) HNRNPU IP-MS samples. Log_2_LFQ enrichment values relative to GFP control are also shown. Proteins with RS domains (curated in (Cascarina and Ross 2022)) are shaded red. **(E, F)** p-values denoting significance of Gene Set Enrichment/Depletion in SAFB and HNRNPU proteomic datasets, corrected for family-wise error rate (FWER; (Olejnik et al. 1997; Subramanian et al. 2005)). -Log10(p-values) for enrichment (EDS sign = 1) and depletion (EDS sign = -1) are shown on the y-axis on positive and negative scales, respectively. **, FWER <0.05.

We selected for further analysis those proteins that were two-fold more abundant (log2 >1) in the SAFB and HNRNPU IPs compared to the GFP IPs. This yielded 69 and 165 proteins that exhibited enriched association with FLAG-SAFB and FLAG-HNRNPU, respectively (Table S6). We next used DAVID to identify enriched Gene Ontology (GO) terms, focusing on the Cellular Component (CC) and Molecular Function (MF) domains (Huang da et al. 2009; Sherman et al. 2022). We observed enrichment of many shared GO terms, including several that center around the themes of RNA-binding and splicing (Figure 5C). Deletion of the DD3 and RGG domains from SAFB and HNRNPU, respectively, led to clear reductions in the association of proteins linked to nucleic acid binding, splicing, and translation (Figure 5C).

We noted that SAFB appeared to associate more robustly with SR proteins than HNRNPU. Conversely, HNRNPU appeared to associate more robustly with other heterogeneous nuclear ribonucleoproteins proteins than SAFB (Figure 5D; Table S6). We also noted a possible enrichment for paraspeckle components in the HNRNPU IPs (Table S6). Therefore, in addition to DAVID analyses, we evaluated separately curated lists of RS-domain proteins (Cascarina and Ross 2022), proteins that biochemically purified along with nuclear speckles/interchromatin granule preparations (Saitoh et al. 2004), and proteins found in paraspeckles (Yamazaki and Hirose 2015). We then determined the relative scale of the enrichment of proteins from each list in SAFB and HNRNPU IPs using an approach modified from GSEA (Subramanian et al. 2005). These analyses showed that nuclear speckle-associated and RS domain-containing proteins were strongly enriched among SAFB interactors (Figure 5E; p-value for enrichment of speckle and RS domain proteins in SAFB, 0.0129 and 0.0038, respectively; Family-wise error rate (FWER) < 0.05 for both tests). Furthermore, these interactions are dependent on the DD3 region of SAFB (Figure 5E). Conversely, HNRNPU interacting proteins were significantly enriched for proteins found in paraspeckles and depleted in association with proteins that harbor RS domains (Figure 5F, p-value for paraspeckle enrichment, 0.0156; p-value for RS domain depletion, 0.0089; FWER < 0.05 for both tests). Thus, SAFB associates with many proteins that co-purify with nuclear speckles, including many that harbor RS domains (Saitoh et al. 2004; Cascarina and Ross 2022), and these associations differ from HNRNPU, another chromatin-associated RNA-binding protein that has previously been implicated in the activation of transcription (Nozawa et al. 2017). These data are consistent with the view that SAFB and HNRNPU promote gene expression through different mechanisms.

## Discussion

SAFB is a multifunctional, chromatin-associated RNA-binding protein essential for mouse development (Garee and Oesterreich 2010; Norman et al. 2016). Using a combination of formaldehyde-based RIP and genetic rescue in ESCs, we observed that SAFB associates primarily but not exclusively with intronic regions of protein-coding genes though purine rich motifs. We also observed that SAFB exhibits a greater-than-expected association with LINE- and Satellite-derived repetitive elements, consistent with prior observations (Aly et al. 2019; Huo et al. 2020). Knockout of SAFB and its paralogue SAFB2 led to differential expression of nearly 1,000 genes associated with multiple biological pathways, including Polycomb targets and developmental regulators. The spliced and unspliced transcripts from the set of genes that were downregulated in DKO ESCs were associated with high levels of SAFB in WT ESCs, consistent with the notion that in these instances the binding of SAFB helps promote overall gene expression. We also found that SAFB associates with many RS domain-containing proteins found in nuclear speckles, as well as with the lncRNA *Malat1*, which is also found in speckles (Hutchinson et al. 2007). The association between SAFB and speckle-associated proteins as well as the ability of a SAFB cDNA to rescue gene expression defects in DKO ESCs each depended on a large, intrinsically disordered domain in SAFB’s C-terminal region which has previously been shown to be important for interaction with RNA and SR proteins (Townson et al. 2004; Finn et al. 2016; Dosztanyi 2018; Meszaros et al. 2018; Corley et al. 2020; Huo et al. 2020). In contrast, HNRNPU, another chromatin-associated RNA-binding protein implicated in transcriptional activation, which under certain assay conditions has also been shown to biochemically co-purify with SAFB (Romig et al. 1992; Renz and Fackelmayer 1996; Nozawa et al. 2017), was more strongly associated with proteins found in paraspeckles, and was depleted in RS domain-containing proteins.

Together, our findings are consistent with a model in which SAFB association with a subset of nascent RNAs promotes overall expression of the corresponding host gene (Figure 6). The exact mechanism by which SAFB association promotes gene expression requires further study. One model involves proteinaceous nuclear condensates called speckles or interchromatin granule clusters. Prior works have demonstrated that SAFB is present in nuclear speckles and associates with other proteins found in speckles (Nayler et al. 1998; Saitoh et al. 2004; Townson et al. 2004). The proteomics data from this study are consistent with those findings. Moreover, it is known that when genes are positioned more closely to nuclear speckles, their transcription is increased (Kim et al. 2020; Zhang et al. 2020). Thus, high levels of SAFB binding to nascent RNA may help position host genes closer to nuclear speckles or speckle-like bodies, which in turn, could promote gene expression at the level of transcription, RNA processing, or both (Figure 6; (Chen and Belmont 2019)). More broadly, our data suggest that the aggregate protein-binding profile of a nascent RNA may modulate the proximity of its host gene to different classes of nuclear condensate (Sabari et al. 2020), a concept that we hope to investigate in the future.

**Figure 6.**
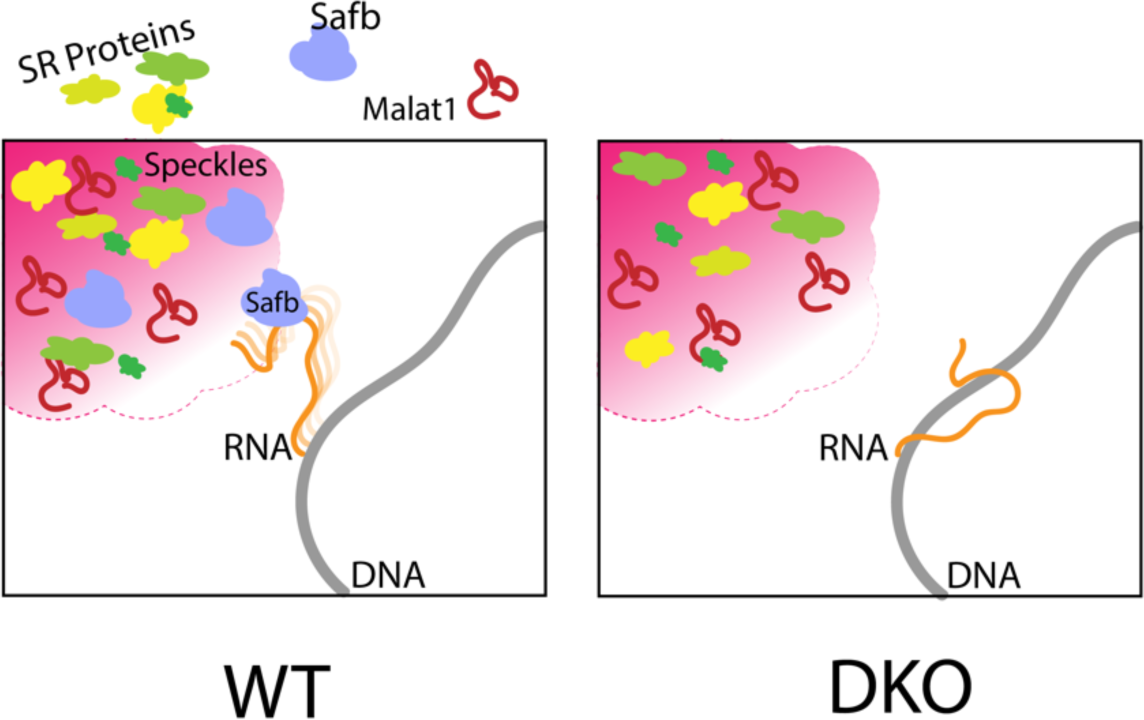
Model: SAFB association with a subset of nascent RNAs may promote overall expression by bringing host genes closer to nuclear speckles, which in turn could promote transcription, splicing, and RNA export (Chen and Belmont 2019).

Only half of the genes whose expression changed significantly upon SAFB/2 DKO experienced downregulation; the other half experienced upregulation. The expression level of many of these upregulated genes shifted back down towards wild-type levels upon reintroduction of SAFB into DKO ESCs, indicating that their dysregulation was reversible and dependent on the continued presence of SAFB. As a class, transcripts from the upregulated set of genes did not associate with high levels of SAFB by RIP, indicating that their increased expression in DKO ESCs is likely due to an indirect effect of SAFB loss.

A significant number of genes whose expression changed upon SAFB/2 DKO and whose expression changes were rescued by reintroduction of SAFB into DKO cells were also altered in analogous fashion upon knockout of the Polycomb protein SUZ12 (Pasini et al. 2007). These included 19 genes whose expression increased upon SUZ12 and SAFB/2 loss, and 85 genes whose expression decreased. Based on these connections, as well as prior studies that have linked SAFB to transcriptional repression and Polycomb-mediated silencing (Townson et al. 2004; Mukhopadhyay et al. 2014; Huo et al. 2020; McCarthy et al. 2021), we profiled H3K27me3 levels by ChIP-Seq H3K27me3 in WT and DKO ESCs (Supplemental Note). We did not observe major changes in the steady-state levels of H3K27me3, *Xist*-deposited H3K27me3, or a connection between the local levels of H3K27me3 and the expression changes induced by SAFB/2 DKO (Supplemental Figure 1). It is possible that the significant overlap between SUZ12 and SAFB/2 dysregulated genes is related simply to the fact that both gene sets were collected from the same cell type -- mouse ESCs. Alternatively, loss of SUZ12 and SAFB/2 may cause some shared changes, for example, to nuclear architecture (Cruz-Molina et al. 2017; Huo et al. 2020), which would then be responsible for the shared changes in gene expression. Note that our conclusions regarding Polycomb are based on H3K27me3 measurements taken at the steady-state, and do not preclude a role for SAFB in establishing new, Polycomb-repressed domains.

In summary, our study provides new insights into the possible regulatory roles of SAFB. In addition to roles in the establishment and maintenance of heterochromatin (Huo et al. 2020; McCarthy et al. 2021), the nuclear retention of RNA (Ron and Ulitsky 2022), and the response to stress (Aly et al. 2019), our work suggests that SAFB may boost the overall expression of certain genes by associating with GA-rich regions in nascent RNA.

## Supplemental Note

### SAFB/2 loss does not cause major disruptions to steady-state levels of H3K27me3 or *Xist*-induced accumulation of H3K27me3

Because of the many links between SAFB and transcriptional repression, including but not limited to PRC2-mediated silencing (Townson et al. 2004; Mukhopadhyay et al. 2014; Huo et al. 2020; McCarthy et al. 2021), because SAFB has previously been shown to associate with the lncRNA *Xist* (Chu et al. 2015; Bousard et al. 2019), and because our own RNA-Seq data show that many of the genes whose expression changes upon SAFB/2 DKO are regulated by PRC2 in ESCs, we asked whether loss of SAFB/2 in ESCs affected H3K27me3 levels. We performed H3K27me3 ChIP-seq in our three WT and two DKO ESC lines, as well as two additional replicate ChIPs for our “Parent WT” and “DKO13” lines. We also performed total H3 ChIP-seq in two replicates each of the Parent WT and DKO13 ESCs as a control to monitor overall H3 density. Using MACS2 (Zhang et al. 2008), we identified a total of 57,909 peaks of H3K27me3 in ESCs: 10,898 Peaks were specific to WT ESCs, 18,837 were specific to DKO ESCs, and 28,174 were detected in both genotypes (Figure S1A). The highest levels of H3K27me3 were found under the peaks that were detected in both WT and DKO cells, and at these peaks, SAFB/2 DKO appeared to result in a slight increase in overall levels of H3K27me3 (Figure S1A, B; Table S7). The WT- and DKO-specific peaks each harbored lower overall levels of H3K27me3 than the set of shared peaks, and likewise exhibited minor differences between genotypes (Figure S1B). In parallel, we examined the levels of H3K27me3 at the promoters of genes whose expression changed upon SAFB/2 DKO and found no major differences between gene classes (Figure S1C). Lastly, our WT and DKO cells are F1-hybrids that express doxycycline-inducible *Xist* from the *Rosa26* locus on the B6-derived copy of chromosome 6 (Trotman et al. 2020). We observed no major differences in the accumulation of H3K27me3 on the *Xist*-expressing allele in WT versus DKO cells (Figure S1D). Together, our data suggest that in mouse ESCs, SAFB does not play a major role in the maintenance of H3K27me3, nor is it required for the deposition of H3K27me3 triggered by expression of *Xist*.

## Methods

### Experimental methods

#### Cell culture

Male mouse ESCs that express doxycycline-inducible *Xist* from the *Rosa26* locus (derivation described in (Trotman et al. 2020)) were grown in: DMEM (Gibco) supplemented with 15% Qualified Fetal Bovine Serum (Gibco), 1% Pen/Strep (Gibco), 1% L-Glutamine (Gibco), 1% Non-Essential Amino Acids (Gibco), 100μM betamercaptoethanol (Sigma) and 0.2% LIF. Cells were maintained in incubators set at 37°C and 5% CO_2_. Media was replaced daily.

#### Generation of WT parent ESCs used for this study

To generate the wild-type parent ESC line from which we ultimately deleted SAFB and SAFB2, full-length *Xist*-expressing ESCs from (Trotman et al. 2020) were deleted of their hygromycin B resistance gene via Lipofectamine transfection of a plasmid expressing FlpE (Addgene #20733; (Beard et al. 2006)). 5μg of FlpE recombinase was mixed with 5μL of P3000 reagent, 7.5μL of Lipofectamine 3000 reagent, and Opti-MEM media (Gibco # 31985-070) to a total volume of 250μL. The reagents were incubated for 5 minutes at room temperature before being added to cells with fresh media. After 24 hours, cells were pulsed with puromycin (2μg/mL) for 72 hours. 96 hours after transfection, ESCs were then trypsinized to single cell suspension and plated onto irradiated fibroblast feeder cells (500-2000 cells / 10cm plate) until individual colonies were visible by eye (4-5 days). Individual colonies were then selected and grown in individual wells for genotyping. After genotyping, candidate clonal colonies underwent hygromycin B (50μg/mL) selection to verify loss of resistance. Genotyping primers used are in Table S8.

#### Generation of SAFB/2 knockout ESCs

sgRNAs to delete SAFB and SAFB2 were designed to the mm10 genome using CRISPOR with the specifications: 20bp-NGG – Sp Cas9, Sp Cas9-Hf1, eSp Cas9 1.1 (Concordet and Haeussler 2018). sgRNA sequences are found in Table S8. Guides were cloned into the pX330 plasmid as specified in ((Cong et al. 2013); Addgene plasmid #42230). To delete SAFB and SAFB2, Parent ESCs were seeded at 0.5×10^6^ cells per well in a 6-well plate. The following day, the cells were transiently transfected using Lipofectamine 3000 (Invitrogen L3000-015): 800ng of sgRNA plasmid pool and 200ng puro resistant GFP plasmid (1μg total) were mixed with 2μL P3000 reagent, 7.5μL Lipofectamine 3000 reagent and with Opti-MEM media (Gibco # 31985-070) to final volume of 250μL. The reagents were incubated for 5 minutes at room temperature before being added to cells with fresh media. After 24 hours, cells were pulsed with Puromycin (2μg/mL) for 48 hours. After puromycin selection, cells were trypsinized to single cells and plated onto irradiated fibroblast feeder cells (500-2000 cells / 10cm plate) until individual colonies were visible by eye (4-5 days). Individual colonies were then selected and grown in individual wells for genotyping.

The two DKO lines that were selected for further study, along with the wild-type Parent line, were then rendered dox-inducible by transfection of the rtTA-expressing plasmid described in (Kirk et al. 2018). One day prior to transfection, parent and DKO ESCs were seeded at 0.5×10^6^ cells per 6-welled well. The following day, 500ng of rtTA plasmid and 500ng of transposase (1μg total DNA) were mixed with 2μL P3000 reagent, 7.5μL Lipofectamine 3000 reagent and with Opti-MEM media (Gibco #31985-070) to 250μL. The reagents incubated for 5 minutes at room temperature before being added to cells with fresh media. After 24 hours, cells underwent G418 selection (50μg/mL) for twelve days.

#### cDNA expression plasmids

Plasmids expressing full-length or truncated versions of SAFB, GFP, and HNRNPU were designed in silico based on existing vector backbones from (Schertzer et al. 2019b), and synthesized by Genewiz. All plasmids are in the process of being deposited into Addgene.

#### Generation of cDNA-expressing ESCs

One day prior to transfection, ESCs were seeded at 0.5×10^6^ cells per well of a 6-well plate. The following day, 850ng of cDNA plasmid and 150ng of piggyBac transposase from (Kirk et al. 2018) were mixed with 2μL P3000 reagent, 7.5μL Lipofectamine 3000 reagent, and with Opti-MEM media (Gibco # 31985-070) to a final volume of 250μL. The reagents were incubated for 5 minutes at room temperature before being added to cells with fresh media. After 24 hours, cells underwent hygromycin B selection (50μg/mL) for one week.

HNRNPU-expressing ESCs were then transfected with the rtTA from (Kirk et al. 2018) as described above. The DKO cells in which GFP and SAFB cDNA constructs were introduced had previously been transduced with rtTA.

#### PCR

Genomic DNA was collected from 0.8×10^6^ cells with 500μL lysis buffer (100mM Tris-HCl pH 8.1, 0.5mM EDTA pH 8.0, 200nM NaCl, 0.2% SDS) + 80μL Proteinase K (Denville) + 8μL linear acrylamide (Thermo Fisher) and incubated at 55°C overnight. Twice the volume of ice cold 100% ethanol was added. Samples were then vortexed and rotated end-over-end at 4°C for 15 min. Samples were spun at max speed for 5 min in at 4°C. The lysis buffer/ethanol mixture was then removed, and the DNA pellet was washed with 70% ethanol, after which the DNA pellet was resuspended in 1x TE (10mM Tris pH 8.0, 1 m EDTA) and incubated overnight at 56°C. DNA concentration was measured via Nanodrop and diluted to 50ng/μL. PCR was performed with ChoiceTAQ (Denville CB4050) as follows: 25μL PCR reaction mixture (2.5μL10x PCR reaction buffer, 0.2μL 10mM dNTPs, 0.25μL 100uM primers, 0.25μL Choice TAQ polymerase, 3μL DNA template (50ng/μL) and ddH2O to 25μL) ran in BioRad C1000 Touch or T100 thermocycler (initial denaturation at 95°C for 3 min; 25 cycles of 95°C for 30s, 58-62.5°C annealing for 30s, and 72°C for 30-45s extension time). PCR primers and conditions are in Table S8.

#### RT-qPCR

Equal amounts of RNA (0.5–1 μg) were reverse transcribed using the High-Capacity cDNA Reverse Transcription Kit (Thermo Fisher Scientific #4368813) with the random primers provided, and then diluted with 30μL 1xTE. For RIP RT-qPCR, 2μL of eluted sample were used in RT reactions. 10μL qPCR reactions were performed using iTaq Universal SYBR Green (Bio-Rad) and custom primers on a Bio-Rad CFX96 system with the following thermocycling parameters: initial denaturation at 95°C for 10 min; 40 cycles of 95°C for 15 s, 60°C for 30 s, and 72°C for 30 s followed by a plate read. The primer concentration used for all qPCR reactions in this study was 0.5μM. Standard curves were used in all qPCR analyses and were prepared by RT of equal volume of wild type sample to other samples. After RT, five 5-fold serial dilutions were made (6 total standards including undiluted RT reaction) and added in duplicate to qPCR plates. After qPCR run, samples were normalized to standard curve read using the BioRad CFX Manager Software. See Table S8 for all primer sequences used.

#### Antibodies

All antibodies used for this study are listed in Table S9.

#### Western blot

To isolate protein for western blotting, 0.8×10e6 cells were washed with 1xPBS and then lysed with 500μL RIPA buffer (10 mM Tris-Cl (pH 7.5), 1 mM EDTA, 0.5 mM EGTA, 1% NP40, 0.1% sodium deoxycholate, 0.1% Sodium Dodecyl Sulfate, 140mM Sodium Chloride) supplemented with 1mM PMSF (Thermo Fisher #36978) and 1x Protease Inhibitor Cocktail (PIC; Sigma Product #P8340). Cell suspensions were rotated for 15 minutes at 4°C, then spun down at high speed at 4°C for 15 minutes and supernatant was collected. Prior to western blotting, protein levels were quantified using the DC assay from Biorad (Product #5000006). 4x SDS loading buffer (Sigma Aldrich Recipe: 0.2 M Tris-HCl pH6.8, 0.4 M DTT, 8% (w/v) SDS, 6 mM Bromophenol blue, 4.3 M Glycerol) was added to samples to 1x final concentration. Samples were then boiled for 5 minutes at 95°C, and equal μg amounts were loaded onto BioRad TGX Stain Free Gels. Samples were run at 50V until past stacking gel, then at 150V for 1-2 hrs. Gels were transferred to PVDF (Immobulon #IPVH00010) membrane either for 1 hour at 125V at 4°C or overnight at 25V at 4°C. Membranes were blocked for 45 minutes in 1xTBST + 5% Milk. Membranes were then incubated with primary antibody either overnight at 4°C or for 1-3 hours at RT. Membranes were washed 3x for 5 minutes each in 1x TBST. Secondary antibodies were diluted in 1xTBST + 5% milk and incubated with membranes for 45 minutes (1:100,000; Invitrogen). Membranes were then washed 3x in 1xTBST washes for 10 minutes, before being imaged with ECL (Thermo Fisher #34096). Antibodies used were FLAG (Sigma F1804, 1:1000), GAPDH (Abcam, ab9484, 1:1000), Total H3 (Proteintech 17168-1-AP, 1:1000), hnRNPU (Santa Cruz sc-32315 1:500), SAFB (Bethyl A300-812A, 1:3000), SAFB2 (Proteintech 11642-1-AP, 1:10,000), V5 (Sigma V8012, 1:1000), Vinculin (Santa Cruz sc-73614, 1:1000), Goat anti-Mouse (Thermo Scientific A16072, 1:100,000) and Goat anti-Rabbit (Thermo Scientific G21234, 1:100,000).

#### Formaldehyde crosslinking of ESCs

For RIP and IP-MS, cells were grown to 75-85% confluency, trypsinized and counted. Cells were washed twice in cold 1xPBS then rotated for 30 min in 10mL of 0.3% formaldehyde (1mL 16% methanol-free formaldehyde (Pierce, #28906) in 49mL 1xPBS) at 4°C. Formaldehyde was quenched with 1mL of 2M glycine for 5 min at room temperature. Cells were washed 3x in cold 1xPBS, then resuspended in 1xPBS at 10×10^6 cells per mL and spun down. PBS was aspirated and pellets were snap frozen in a liquid nitrogen bath and immediately transferred to -80°C until further processing.

For ChIP, cells were grown to 75-85% confluency and counted. Cells were washed once with 1xPBS and crosslinked with 0.6% formaldehyde for 10 minutes at room temperature. Formaldehyde was quenched with 557 μL of 2.5 M glycine and washed twice with cold 1xPBS. Cells were then scraped in 2mL cold 1xPBS supplemented with 1x Protease Inhibitor Cocktail (PIC; Sigma Product #P8340). 10mLs 1xPBS + 0.05% tween-20 were added to collect the cells. Cells were spun down, resuspended in 1xPBS at 10×10^6 cells per mL and spun down. PBS was aspirated and pellets were snap frozen in a liquid nitrogen bath and immediately transferred to - 80°C for storage.

#### RNA-IPs (RIPs)

RIPs were performed similar to (Schertzer et al. 2019a), which is a protocol originally adapted from (Hendrickson et al. 2016; Raab et al. 2019). 25μL protein A/G agarose beads (Santa Cruz sc-2003) were washed three times in blocking buffer (0.5% BSA in 1xPBS) and incubated overnight at 4°C with 10μL antibody (anti-SAFB; Bethyl 812-300A; FLAG, Sigma F1804; V5, Sigma V8012) or 10μg of mouse IgG (Invitrogen). 10×10^6 cells were resuspended in 500μL RIPA Buffer (50mM Tris-HCl, pH8, 1% Triton X-100, 0.5% sodium deoxycholate, 0.1% SDS, 5mM EDTA, 150mM KCl) supplemented with 1x Protease Inhibitor Cocktail (PIC; Sigma Product #P8340), 2.5μL SuperaseIN (Thermo Fisher Scientific AM2696) and 0.5mM DTT and sonicated twice for 30s on and 1 min off at 30% output using the Sonics Vibracell Sonicator (Model VCX130, Serial# 52223R). Samples were spun down at high speed and 50μL total lysate was saved for input. Beads were washed three times in 1mL fRIP buffer (25mM Tris-HCl pH 7.5, 5mM EDTA, 0.5% NP-40, 150mM KCl) and resuspended in 450μl fRIP buffer supplemented as above with PIC, SuperaseIN and DTT, then mixed with sonicated samples. Samples were rotated overnight at 4°C, then washed once with 1mL fRIP buffer and resuspended in 1mL PolII ChIP Buffer (50mM Tris-HCl pH 7.5, 140mM NaCl, 1mM EDTA, 1mM EGTA, 1% Triton X-100, 0.1% Sodium-deoxycholate, 0.1% Sodium dodecyl sulfide) before transferring to a new 1.7mL tube. Samples were rotated at 4°C for five minutes, spun down at 1200xg, and the supernatant aspirated. Samples were washed twice more with 1mL PolII ChIP Buffer, twice with 1mL High Salt ChIP Buffer (50mM Tris-HCl pH 7.5, 500mM NaCl, 1mM EDTA, 1mM EGTA, 0.1% sodium-deoxycholate, 0.1% sodium dodecyl sulfide, 1% Triton X-100), and once in 1mL LiCl buffer (20mM Tris pH 8.0, 1mM EDTA, 250mM LiCl, 0.5% NP-40, 0.5% sodium-deoxycholate); each wash included a five-minute rotation at 4°C. At the final wash, samples were transferred to a new 1.7mL tube. After the final wash, inputs were thawed on ice and bead samples were resuspended in 56μL water, 33μL of 3x reverse-crosslinking buffer [3x PBS, 6% N-lauroyl sarcosine and 30mM EDTA], 5μL 100mM DTT, 20μL Proteinase K, and 1μL of SuperaseIN. Samples were incubated for 1hr at 42°C, then 1hr at 55°C, then 65°C for 30 minutes, and mixed by pipetting every 15 minutes. Afterwards, 1mL Trizol was added, samples were vortexed, 200μL CHCl3 was added, samples were vortexed, and finally spun at 12000xg for 15 mins at 4°C. The aqueous phase was then extracted and to that one volume of 100% ethanol was added. Samples were vortexed and applied to Zymo-Spin IC Columns (from #R1013) and spun for 30 seconds at top speed on a benchtop microcentrifuge. 400μL of RNA Wash Buffer (Zymo #R1013) was added and samples were spun at top speed for 30 seconds. For each sample, 5μL DNase I and 35μL of DNA Digestion Buffer (Zymo #R1013) was added directly to the column matrix and incubated at room temp for 20 minutes. 400μL of RNA Prep Buffer was then added (Zymo #R1013), and columns were spun at top speed for 30 seconds. 700μL RNA Wash Buffer (Zymo #R1013) was then added, and columns were spun at top speed for 30 seconds. 400μL RNA Wash Buffer was then added, and columns were spun at top speed for 30 seconds. The flow through was discarded and columns spun again for 2 minutes to remove all traces of wash buffer. Columns were transferred to a clean 1.7mL tube, 15μL of ddH2O was added to each column, and after a five-minute incubation, samples were spun at top speed to elute.

#### Total RNA isolation

ESCs were grown in 6-well plates to ∼80% confluency. Cells were washed twice with 1x PBS and 1 mL of TRIzol was added per well. Samples were pipetted up and down at least 10 times, transferred to a microcentrifuge tube and briefly vortexed. Samples were incubated at RT for 5 minutes then 200μl of chloroform was added. Afterwards, samples were vortexed and incubated for 3 minutes at RT. Samples were spun down at 12000xg for 15 min at 4°C. The upper aqueous phase was moved to a new tube and 8μL of linear acrylamide (Thermo Fisher, AM9520) was added. Then 500μL of 100% isopropanol was added, and samples were vortexed and incubated at RT for 10 minutes. Tubes were spun down 12000xg for 10 minutes at 4°C. Supernatant was removed and pellets washed with 1mL of cold 75% ethanol. Samples were briefly vortexed and spun down at 7500xg for 5 min at 4°C. Supernatant was discarded and pellet was dried by repeated spin down and aspiration. Final pellets were resuspended in 100μL water by pipetting up and down.

#### RNA sequencing

For RIP-Seq inputs, 100ng of RNA prepared from RIP input samples was used for library preparations. For RIP-Seq RIP samples, 9μL of RIP sample (from 15μL total) were used. For total RNA-Seq, 900ng of total RNA was used. Each library preparation included 1μL of 1:250 dilution of ERCC Spike-In RNAs (Ambion #4456653). 10μL total were prepped using the KAPA RNA HyperPrep Kit with RiboErase (Kapa Biosystems; product #KR1351). Sequencing was performed on an Illumina Next-Seq 500, using high-output, 75-cycle kits.

#### H3K27me3 and total H3 ChIP-Seq

The day before sonication, 25μL of protein A/G agarose beads (Santa Cruz sc-2003) were washed 3 times in block solution (0.5% BSA in 1xPBS) before being resuspended in 300μL blocking solution. 10μL per 10 million cells of antibody (Abcam mouse monoclonal ab6002) was added, then beads and antibody conjugated via rotation overnight at 4°C.

On the day of sonication, 10 million ESCs crosslinked with 0.6% formaldehyde were re-suspended in lysis buffer 1 (50mM HEPES pH 7.3, 140mM NaCl, 1mM EDTA, 10% glycerol, 0.5% NP-40, 0.25% Triton X-100, and 1x PIC (PIC; Sigma Product #P8340) incubated for 10 minutes at 4°C, and then incubated with lysis buffer 2 (10mM Tris-HCl pH 8.0, 200mM NaCl, 1mM EDTA pH 8.0, 0.5 mM EGTA pH 8.0, and 1x PIC) for 10 minutes at room temperature. For H3K27me3 ChIPs, cells were re-suspended in lysis buffer 3 (10 mM Tris-HCl pH 8.0, 100 mM NaCl, 1 mM EDTA pH8.0, 0.5 mM EGTA pH 8.0, 0.1% Na-deoxycholate, 0.5% N-lauroyl sarcosine, and 1x PIC) and then sonicated.

ChIPs were then performed by incubating sonicated cell lysates at a concentration of 20 million cells/1ml of lysis buffer 3 containing 1% Triton X-100 with pre-conjugated with H3K27me3 antibody/agarose beads overnight at 4°C. After overnight H3K27me3 ChIP, beads were washed 5x in RIPA buffer (50 mM HEPES pH 7.3, 500 mM LiCl, 1 mM EDTA, 1% NP-40 and 0.7% Na-Deoxycholate) for 5 minutes each and then once in TE. To elute the DNA, beads were re-suspended in Elution buffer (50mM Tris pH 8.0, 10mM EDTA, and 1% SDS) and placed on a 65°C heat block for 17 minutes with frequent vortexing. Crosslinks were reversed overnight at 65°C, eluates were incubated with Proteinase K and RNase A, and DNA was extracted with phenol/chloroform and precipitated with ethanol. DNA was prepared for sequencing on the Illumina platform using Next Reagents (NEB) and Agencourt AMPure XP beads (Beckman Coulter).

#### IP-mass spectrometry sample preparation

40μL protein A/G agarose beads (Santa Cruz sc-2003) were washed three times in blocking buffer (0.5% BSA in 1xPBS) and incubated overnight at 4°C with 20μL antibody (anti-FLAG; Sigma F1804-1MG). 30×10^6 of ESCs crosslinked with formaldehyde as described above were resuspended in 500μL RIPA Buffer (50mM Tris-HCl, pH8, 1% Triton X-100, 0.5% sodium deoxycholate, 0.1% SDS, 5mM EDTA, 150mM KCl) supplemented with (1x PIC (PIC; Sigma Product #P8340), 2.5μL SuperaseIN (Thermo Fisher Scientific product #AM2696), and 0.5mM DTT) and sonicated twice for 30 s on and 1 min off at 30% output using Sonics Vibracell Sonicator (Model VCX130, Serial# 52223R). Samples were spun down at high speed and 50μL total lysate was saved for input. Beads were washed 3 times in fRIP buffer (25mM Tris-HCl pH 7.5, 5mM EDTA, 0.5% NP-40, 150mM KCl) and resuspended in 450μL fRIP buffer supplemented with 1x PIC (PIC; Sigma Product #P8340), 2.5μL SuperaseIN (Thermo Fisher Scientific product # AM2696) and 0.5mM DTT to bring samples to a 1:1 ratio of RIPA/fRIP buffer. Samples rotated overnight at 4C. At 4C, samples were then washed once with 1mL fRIP buffer and then resuspended in 1mL PolII ChIP Buffer (50 mM Tris-HCl pH 7.5, 140 mM NaCl, 1 mM EDTA, 1 mM EGTA, 1% Triton X-100, 0.1% Na-deoxycholate, 0.1% SDS) and transferred to a new Eppendorf tube. Samples were washed two more times with PolII ChIP Buffer, twice with High Salt CLIP Buffer (50 mM Tris-HCl pH 7.4, 1 M NaCl, 1 mM EDTA, 1% NP-40, 0.5% Na-deoxycholate, 0.1% SDS), and resuspended in 1mL LiCl buffer (20 mM Tris pH 8.0, 1 mM EDTA, 250 mM LiCl, 0.5% NP-40, 0.5% Na-deoxycholate) and moved to a new microcentrifuge tube. Samples were then resuspended in cold 1xPBS and moved to a new Eppendorf tube. Samples were washed 3x with 1mL cold 1xPBS. 25% of samples were saved for western blot, and the remaining 75% were subjected to on-bead trypsin digestion as previously described (Rank et al. 2021). Briefly, after the last wash buffer step during affinity purification, beads were resuspended in 50μL of 50mM ammonium bicarbonate (pH 8). On-bead digestion was performed by adding 50μL 50mM ammonium bicarbonate (pH8) and 1μg trypsin and incubated, shaking, overnight at 37°C. Beads were pelleted and transferred supernatants to fresh tubes. The beads were washed twice with 100μL LC-MS grade water, and washes added to the original supernatants. Samples were acidified by adding formic acid to final concentration of 2%, to pH ∼2. Peptides were desalted using peptide desalting spin columns (Thermo), lyophilized, and stored at -80°C until further analysis.

#### LC/MS/MS analysis

The peptide samples were analyzed by LC/MS/MS using an Easy nLC 1200 coupled to a QExactive HF Biopharma mass spectrometer (Thermo Scientific). Samples were injected onto an Easy Spray PepMap C18 column (75μm id × 25 cm, 2μm particle size) (Thermo Scientific) and separated over a 2 hr method. The gradient for separation consisted of 5–45% mobile phase B at a 250 nl/min flow rate, where mobile phase A was 0.1% formic acid in water and mobile phase B consisted of 0.1% formic acid in acetonitrile (ACN). The QExactive HF was operated in data-dependent mode where the 15 most intense precursors were selected for subsequent fragmentation. Resolution for the precursor scan (m/z 350–1700) was set to 60,000, while MS/MS scans resolution was set to 15,000. The normalized collision energy was set to 27% for HCD. Peptide match was set to preferred, and precursors with unknown charge or a charge state of 1 and ≥ 7 were excluded.

### Computational analyses

#### RIP- and RNA-Seq alignment

RIP and RNA-Seq data were aligned to the mm10 mouse genome using STAR with default parameters (Dobin et al. 2013). Alignments with a quality score ≥30 were retained for subsequent analyses (Li et al. 2009). For analysis of multimapping reads with TElocal (Jin et al. 2015), additional parameters (--winAnchorMultimapNmax 100 and --outFilterMultimapNmax 100) were specified in STAR alignments.

#### RIP peak calling

After alignment and filtering, all SAFB RIP-Seq data from WT ESC replicates were concatenated, and using samtools, were split into two files, corresponding to alignments that mapped to the positive and negative strands of the genome, respectively. Using a custom perl script, the strand information within the positive and negative strand alignment files was randomized so as to better match the criteria of the MACS peak caller, which uses the average distance between positive and negative strand alignments to estimate the fragment length (Zhang et al. 2008). Putative peaks were called on strand-randomized positive and negative strand alignment files, respectively, using default MACS parameters and not providing a background file (Zhang et al. 2008). Peak bed files were converted to SAF format and reads under each putative peak were counted from SAFB RIP-Seq alignments performed in WT and DKO ESCs using featureCounts (Liao et al. 2014). We retained putative peaks that were represented by at least five reads in at least two of the three WT ESC lines profiled. We then used DESeq2 under default parameters to identify those putative peaks that were ascribed a p-value of <0.05 by DESeq2 when comparing signal between SAFB/2 DKO ESCs and wild-type ESCs. Lastly, we retained only those putative peaks that harbored an average aligned-reads-per-million-total-reads (RPM) signal of at least two-fold less in SAFB/2 DKO ESCs compared to wild-type ESCs. This yielded 32,354 regions that were potentially enriched in their association with SAFB in wild-type ESCs. As a final filtering step, we used featureCounts to count the number of reads under these 32,354 regions that aligned to the mm10 genome with quality scores of ≥30 from within the SAFB-FLAG and GFP-FLAG RIP-Seq datasets. From the initial set of 32,354 regions, 23,853 had a total of at least 5 reads distributed between the SAFB-FLAG and GFP-FLAG; of these, 1,356 regions had higher signal in the GFP-FLAG compared to the SAFB-FLAG RIP-Seq dataset and were dropped from further analysis, yielding a total of 30,998 regions that we defined as SAFB-associated peaks (Table S1).

#### RIP scatter plots

Scatter plots in Figure 2 were constructed using featureCounts to count the reads under each of the 30,998 SAFB peaks in each dataset. Read counts were then plotted using R (Team 2017).

#### SAFB motif analysis

To identify the motifs associated with SAFB peaks, we provided the sequences of the three thousand peaks with the greatest level of SAFB signal (top ∼10% of peaks) as input to the Sensitive, Thorough, Rapid, Enriched Motif Elicitation tool (STREME) from the MEME Suite (Bailey et al. 2015; Bailey 2021). Randomized control sequences with lengths the same as each of the peak sequences were developed with weighted nucleotide occurrence based on the mononucleotide content of the mm10 reference genome. The --rna flag was specified to account only for single-stranded analysis and motif width was restricted to between four and eight nucleotides; the motifs with the top three most significant p-values are shown in Figure 2D.

#### UCSC wiggle density plots

UCSC wiggle density plots were made from filtered sam files using custom perl scripts.

Tracks of individual and pooled replicates are located here: https://genome.ucsc.edu/s/recherney/Cherney_Safb_2022

#### Intersection of SAFB peaks and genic features

To identify the genic features under each SAFB peak, the GENCODE Basic vM25 GTF was downloaded and modified to include annotations of 5’ and 3’ UTRs (Frankish et al. 2021). From this file, features mapping to protein-coding genes (GENCODE transcript_type: “protein_coding”) and lncRNAs (GENCODE transcript_types: “bidirectional_promoter_lncRNA”, “macro_lncRNA”, “antisense”, “3prime_overlapping_ncRNA”, “lincRNA”, “processed_transcript”, “sense_intronic”, and “sense_overlapping”) were extracted and intersected with SAFB peaks using bedtools (Quinlan and Hall 2010). Peaks were classified as exon-overlapping if they fell within a gene and overlapped >50% of the exon in question, otherwise they were classified as intron-overlapping. The classification of each peak can be found in Table S1.

#### Intersection of SAFB peaks and repeatmasked elements

To determine whether SAFB peaks overlapped with repeat-masked genomic elements more than would be expected by random chance, repeat-masked elements were first extracted from all 20 mouse autosomes and the X chromosome using the UCSC genome browser MySQL relational database (Lee et al. 2022). Peaks were then intersected with repeat-masked elements using bedtools (Quinlan and Hall 2010). To estimate what level of intersection might be expected from random chance, we performed 1,000 repetitions of the following process: the starting position of each peak in the set of SAFB peaks was shifted randomly to a new position between 2000 and 10000 bases upstream or downstream; then, each complete set of randomized set of peaks was intersected with repeat-masked elements extracted from UCSC. The error bars in Figure 2G represent the standard deviation of the number of intersections with each class of repeat-masked element from each set of randomized peaks.

#### Intersection of multimapping RIP reads with repeat-derived elements

To determine the relative representation of repeat-derived elements in multimapping reads from SAFB RIPs, reads were aligned to mm10 with STAR using the parameters recommended by TElocal (--winAnchorMultimapNmax 100 and --outFilterMultimapNmax 100; (Jin et al. 2015)). samtools was then used to extract alignments with MAPQ = 0 (i.e., multimapping reads; (Li et al. 2009)). Relative read abundance over repeat-derived elements was then calculated with TElocal, using the pre-built ‘mm10_rmsk_TE.gtf.locInd’ index and the ‘--stranded revers’ option (Jin et al. 2015). Counts from TElocal were converted to RPM values (reads-per-million-total-reads). We retained only those elements that were represented by an RPM value of ≥1 summed across all datasets, were represented by an average RPM of ≥0.25 in the WT RIPs, and that had ≥2-fold higher RPM in the WT compared to DKO RIPs. To determine the expected representation of each class of repeat in this final list of filtered elements, we summed the genomic space occupied by each class of repeat in the mm10_rmsk_TE.gtf from TElocal, and used this information to calculate the expected genomic space occupied by each class of repeat in our final list of filtered elements. We then used Chi-squared tests to determine whether the actual genomic space occupied by each class of repeat in our final list of filtered elements differed significantly from what was expected. Classes of repeat that were represented by less than 25 elements in our final filtered list were not plotted in Figure 2H. Processed data used to generate Figure 2H are included in Table S2.

#### Differential gene expression analyses

To detect genes that were differentially expressed between WT and SAFB/2 DKO ESCs, we performed RNA-Seq on “Input” RNA extracted from the same sonicated extracts of formaldehyde-crosslinked WT and DKO ESCs samples that were used to perform SAFB RIP-Seq described in Figure 2 – the RNA extraction protocol is detailed in the “*RNA-IPs*” section of the methods. “Input” RNA Reads were aligned to mm10 using STAR and default parameters (Dobin et al. 2013), alignments were filtered to retain only those reads with a mapping-quality of ≥30 using samtools (Li et al. 2009), and then the number of filtered reads mapping to each GENCODE vM25 gene was counted using featureCounts (Liao et al. 2014): [-g gene_name -s 2 -a gencode.vM25.basic.annotation.gtf -o]. Genes that had less than 10 total reads summed across all five samples (three WT and two DKO) were excluded from downstream analyses. Counts were loaded into DESeq2, and the genes that had adjusted p-values for differential expression between WT and DKO samples of <0.05 were retained and reported as “significant” in Table S3.

#### Gene set enrichment analyses

Gene set enrichment analyses of differentially expressed genes were performed using the Molecular Signatures Database webserver (Liberzon et al. 2015). The lists of significantly down- and upregulated genes were input, orthology mapped onto the human genome, and queried for overlap with the Hallmark, Chemical and Genetic Perturbations, and GO Biological Process Gene Sets. The top 20 most significantly-overlapping gene sets from each search are reported in Table S4.

#### Assessing SAFB RIP signal over spliced and unspliced transcripts

To determine the extent to which SAFB was enriched over expressed spliced and unspliced transcripts, we took advantage of the kallisto algorithm, which was designed to enable probabilistic alignment of short-read RNA-Seq data (Bray et al. 2016). To enable detection of unspliced transcripts, we created a version of the GENCODE vM25 basic transcriptome that for each gene, included one representative unspliced transcript that began at the first annotated transcription start and the last annotated transcription end (vM25_basic_complete.fa).

In parallel, because like all RIP- or CLIP-Seq datasets, our RIP-Seq datasets contained reads that align to genomic regions that were not classified as peaks, we selected for our downstream analyses only the subset of RIP-Seq reads that aligned under SAFB peaks. To do this, SAFB RIP-Seq reads from WT and DKO datasets were aligned to mm10 using STAR and filtered for mapping quality ≥30 using samtools (Li et al. 2009; Dobin et al. 2013). We then used samtools to split the alignments by strand, and for each stranded alignment file, again using samtools, selected the subset of alignments from the WT and DKO datasets that aligned under each SAFB peak. Still using samtools, we converted the bam alignments back into fastq data. These final fastq files represent the subset of RIP-Seq data that aligned under each classified peak and exclude most noise in the WT dataset. Input RNA-Seq and the subset RIP-Seq data were then aligned with kallisto to an index made from vM25_basic_complete.fa, using the options [-l 200 -s 50 --rf-stranded]. For each transcript isoform in vM25_basic_complete.fa, TPM counts reported from the DKO RIP-Seq dataset were subtracted from TPM counts in the WT RIP-Seq dataset.

The output file then underwent the following filtering parameters: (1) Transcripts isoforms that we had previously filtered out prior to performing DESeq analyses were excluded, (2) For each gene remaining, we retained the single spliced isoform with the highest expression level as representative, and (3) Transcript isoforms whose expression in the WT input total RNA-Seq data were less than 0.125 TPM were excluded, including 25 and 14 genes that were originally called significantly up- and downregulated by DESeq2, respectively. Finally, we split the transcripts of the genes that did not significantly change in expression upon DKO into three categories: those with low (0.125 -1 TPM), medium (>1 but <16 TPM) and high (>16 TPM) levels of expression. Data used to make plots in Figure 3D are included in Table S5.

#### IUPred2

Protein disorder plots for SAFB were constructed using the webserver for IUPred2A (Erdos and Dosztanyi 2020).

#### Kallisto analyses of rescue RNA-Seq data

To determine the relative abundance of spliced and unspliced transcript isoforms between the SAFB, SAFB-*Δ*DD3, and GFP rescue datasets, RNA-Seq data were aligned using kallisto to an index made from vM25_basic_complete.fa, using the options [-l 200 -s 50 --rf-stranded]. The output file then underwent the same filtering as in Figure 3D: (1) Transcripts isoforms that we had previously filtered out prior to performing DESeq analyses were excluded (62,934), (2) For each gene remaining, we retained the single spliced isoform with the highest expression level as representative, and lastly, (3) Transcript isoforms whose averaged expression of the two SAFB-FL-WT replicates in the total RNA-Seq data were less than 0.125 TPM were excluded, including 23 and 20 genes that were originally called significantly up- and downregulated by DESeq2, respectively. We then normalized each individual replicate TPM value to the SAFB-FL-WT TPM average. P-values were determined using a paired t-test. Data used to make plots in Figure 4D and E are included in Table S5.

#### Mass spectrometry data analysis

Raw data files were processed using MaxQuant version 1.6.15.0 and searched against the reviewed mouse database (containing 17,051 entries), appended with a contaminants database, using Andromeda within MaxQuant. Enzyme specificity was set to trypsin, up to two missed cleavage sites were allowed, and methionine oxidation and N-terminus acetylation were set as variable modifications. A 1% FDR was used to filter all data. Match between runs was enabled (5 min match time window, 20 min alignment window), and a minimum of two unique peptides was required for label-free quantitation using the LFQ intensities. Perseus was used for further processing (Tyanova et al. 2016). Only proteins with >1 unique+razor peptide were used for LFQ analysis. Proteins with 50% missing values were removed and missing values were imputed from normal distribution within Perseus. Log2 fold change (FC) ratios were calculated using the averaged Log2 LFQ intensities of IP sample compared to GFP control. Proteins with Log2 FC >1 were considered biological interactors and analyzed further. All analyzed protein interaction data are present in Table S6. The mass spectrometry proteomics data have been deposited to the ProteomeXchange Consortium via the PRIDE partner repository (Perez-Riverol et al. 2022) with the dataset identifier PXD038103. Reviewer account details: [Username; Password: 7TiboSU8].

#### DAVID GO analysis

Gene Ontology analyses were conducted using DAVID (Huang da et al. 2009; Sherman et al. 2022). Genes were searched against UP_KW_BIOLOGICAL_PROCESS, UP_KW_CELLULAR_COMPONENT, UP_KW_MOLECULAR_FUNCTION, GOTERM_BP_DIRECT, GOTERM_CC_DIRECT and GOTERM_MF_DIRECT. The list shown in Figure 5 represents the union of the top twelve most enriched CC and MF GO terms from the wild-type SAFB and HNRNPU pulldowns that also passed an FDR of <0.01. The bubbleplot in Figure 5 was created using tidyverse v 1.3.1 package in R version 4.0.4 (Team 2017; Wickham et al. 2019).

#### Custom gene set enrichment analyses

To determine whether the SAFB and HNRNPU immunoprecipitates were enriched in proteins found in nuclear speckles (Saitoh et al. 2004), paraspeckles (Yamazaki and Hirose 2015), or proteins that harbor RS domains (Cascarina and Ross 2022), we used the lists of proteins reported in the aforementioned references and the 69 and 165 proteins that we classified as enriched over GFP control in the SAFB and HNRNPU IPs, respectively (Table S6). We followed the gene ranking metric referred to as ‘log2 Ratio of Classes’ (LRC) and the Gene Set Enrichment Analysis (GSEA) framework described in (Subramanian et al. 2005). However, we decided to use a custom version of GSEA, instead of the standard version, to account for i) the limited number of genes in our two datasets, ii) the number of replicates available (two for each sample), which is lower than the canonical threshold of at least 7 recommended by the GSEA authors for phenotype permutations, and iii) the few gene sets (only three), which were tested at the same time. Briefly, prior to calculating enrichments, the average LFQ values per protein per dataset were calculated and divided by the average LFQ values of each protein in the GFP IP; this ratio was then log2 transformed. Enrichment and depletion scores were then calculated separately for each dataset. To calculate enrichment/depletion score (EDS), for each gene set and dataset of interest, we first converted log2-transformed ratios into a ranked list. The highest rank was defined as the numerical value that corresponds to the total number of rows in the list in question and was assigned to the corresponding gene in the dataset that had the highest log2-transformed ratio in the list. The lowest rank was defined as a value of 1 and was assigned to the gene in the dataset that had the lowest log2-transformed ratio in the list. These ranks were then assigned to the genes of each gene set and their averages became the EDSs, which are specific for each dataset (SAFB and HNRNPU), gene set (speckle, paraspeckle and SR proteins) and condition (full protein vs. protein with a deleted domain). The neutral point (NP) for each dataset in each gene set is equal to the [(#genes in the dataset +1)/2]. Specifically, the EDS of each gene set was then defined as the average rank of genes in each dataset that were present in the gene set. Gene sets whose EDS > NP were classified as ‘enriched’ and those whose EDS < NP were classified as depleted. To assess statistical significance, we generated random EDS values by averaging the ranks of as many randomly-selected dataset genes as those present in each gene set and repeated this process, for each dataset, each gene set and each protein form (full or with a deleted domain) 100,000 times. Then, we assessed the probability that each EDS was produced by chance following the approach outlined in (Mielke and Berry 2007), in which the p-value for enrichment ≈ [(number of permutated cases with EDS ≥ NP)/(number of total permutations performed)] or, in the case of depletion, using as numerator of this ratio [# permutated gene sets with EDS ≤ NP]. Resulting p-values were Bonferroni-corrected, thus controlling for the family-wise error rate (FWER) as recommended in (Olejnik et al. 1997; Subramanian et al. 2005). The FWER was assessed at three levels: 0.10 (*), 0.05 (**) and 0.01 (***). We performed Bonferroni correction by keeping together enriched and depleted gene sets, when they were present at the same time (namely, in the HNRNPU dataset of the full protein), thus producing more conservative statistical results than performing the correction after splitting enrichment and depletion cases, as done in standard GSEA.

#### Analysis of H3K27me3 ChIP-Seq data

H3K27me3 data were merged by genotype (WT and DKO, respectively) and total H3 data were merged. Data were then aligned to mm10 using bowtie2 (Langmead and Salzberg 2012). Peaks were called using MACS2 with total H3 as a control under the following parameters: [macs2 callpeak -t -c bam -n -f BAM -g mm –broad –broad-cutoff 0.01] (Zhang et al. 2008). H3K27me3 peak locations are included in Table S7. WT and DKO peak files were catenated and then merged using bedtools [bedtools merge -I in.file > out.file]. The independent WT and DKO data sets were intersected with the merged data file [bedtools intersect -wao -header -a union file -b wt or dko file > outfile] to identify WT and DKO specific H3K27me3 peaks.

#### H3K27me3 analyses

To annotate gene promoters, we took the gencode.vM25.basic.annotation.gtf file and filtered features for transcripts only. We then took the coordinate of the transcription start site from each transcript and added 2kb upstream and 1kb downstream (3kb total) and used this region as the transcript promotor region. We then used featureCounts (featureCounts -s 0 -F SAF -a promoter.file -o out.file) to align H3K27me3 reads to our annotated promoters. We then used bedtools (bedtools intersect -wao -header -a promoter.file -b k27peaksfile > out.file) to intersect our previously called H3K27me3 peaks in our WT and DKO files with our promoter file to find the levels of H3K27me3 at promoters.

To perform allelic H3K27me3 analyses, variant sequence data for the mm10 genome build was obtained from the Sanger Institute (http://www.sanger.ac.uk/resources/mouse/genomes/; (Keane et al. 2011), and a CAST/EiJ (CAST) pseudogenome was created as in (Calabrese et al. 2012; Calabrese et al. 2015). Reads were aligned to both the B6 and CAST version of mm10 using STAR, and those that had a mapping quality greater than or equal to 30 were extracted with samtools. Reads that overlapped B6 or CAST SNPs were detected using a custom perl script as in (Calabrese et al. 2012; Calabrese et al. 2015). Allelic tiling density plots over chr6 were created as in (Schertzer et al. 2019a), using a bin size of 4,000bp.

## Acknowledgements

We thank UNC colleagues, particularly Daniel Dominguez, for many helpful discussions.

## Figures

**Figure S1.**
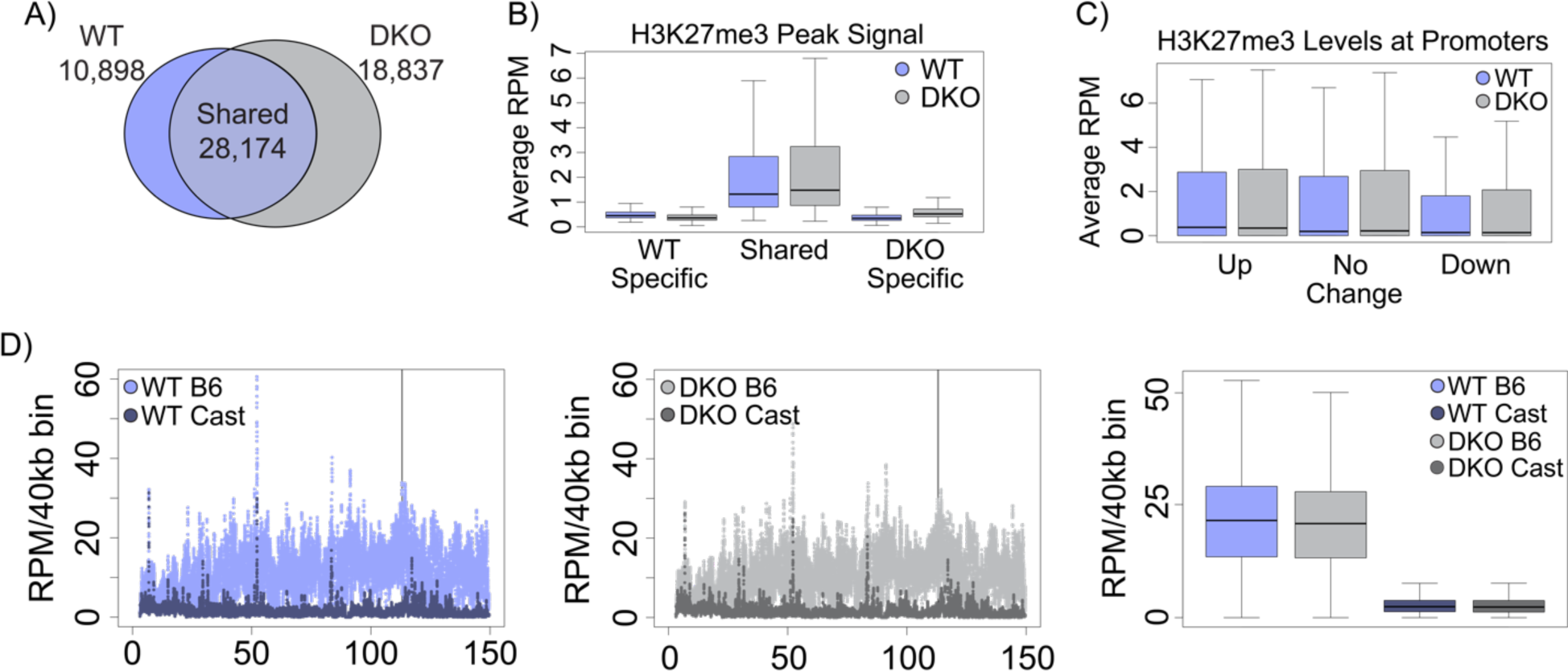
SAFB/2 loss does not cause major disruptions to steady-state levels of H3K27me3 or *Xist*-induced accumulation of H3K27me3. **(A)** The numbers of H3K27me3 peaks specific to WT or DKO ESCs, or shared between both genotypes. **(B)** Average RPM of H3K27me3 signal under genotype-specific and shared H3K27me3 peaks. **(C)** H3K27me3 levels at promoters of genes that are significantly up or down regulated, or do not change in expression, upon SAFB/2 DKO. **(D)** Allelic H3K27me3 levels on chr6 in WT and DKO ESCs. *Xist* is expressed from the B6 allele. Boxplots summarizing allelic H3K27me3 levels on chr6 are also shown.

